# Effects of *P16* DNA Methylation on Proliferation, Senescence, and Lifespan of Human Fibroblasts

**DOI:** 10.1101/405407

**Authors:** Ying Gan, Chenghua Cui, Shengyan Xiang, Baozhen Zhang, Dajun Deng

## Abstract

The aim is to study the effects of *P16* DNA methylation on lifespan of normal cells. An expression-controllable pTRIPZ vector expressing P26-specific zinc finger binding protein-based methyltransferase (P16-Dnmt) was used to induce *P16* methylation in primary CCD-I8C0 fibroblasts via stable transfection. Long-term dynamic IncuCyte analysis showed that CCD-I8C0 fibroblasts expressing baseline P16-Dnmt continued proliferating until passage-26 in the 53^th^ post-transfection week, while vector control cells stopped proliferating at passage-6 and completely died 2 weeks later. The proliferation rate of baseline P16-Dnmt cells was significantly higher than that of vector control cells. The proportion of β-galactosidase-positive staining cells was significantly decreased in baseline P16-Dnmt cells compared to vector control cells. The P16 expression was lost in baseline P16-Dnmt cells at and after passage-6. The average telomere length in baseline P16-Dnmt cells also gradually decreased. In conclusion, *P16* methylation could prevent senescence, promote proliferation, and expand lifespan of human fibroblasts, which may play a role in cancer development.

**Summary:** A zinc finger protein-based DNA methyltransferase (P16-Dnmt) expressed at the baseline level could specifically methylate *P16* promoter CpG islands. *P16* methylation induced by baseline P16-Dnmt could significantly prevent senescence, promote proliferation, and expand lifespan of primary human fibroblasts.

## INTRODUCTION

*P16* gene *(CDKN2A)* is most frequently inactivated in cancer genomes by allele deletion and methylation of CpG islands around transcription start site (TSS) (1-5). *P16* germline inactivation causes high susceptibility to familial melanoma and pancreatic cancer (6,7). Abnormal *P16* methylation is prevalent in cancer and precancerous tissues (5). *P16* methylation inactivates transcription of genes, including *ANRIL* located within the *CDKN2A/B* gene antisense-strand (8,9). Many studies showed that *P16* methylation significantly increased risk of malignant transformation of precancerous tissues such as epithelial dysplasia and Barrett’s esophagus (10-15). These phenomenons imply that *P16* methylation maybe a driver in cancer development. However, solid evidence to prove that epigenetic *P16* inactivation drives malignant transformation of normal cells is still needed.

Recently, epigenetic editing tools with different editing efficiencies and specificities are emerging which are based on engineered ZFP (zinc finger protein), CRISPR (clustered regularly interspaced short palindromic repeats)/dCas9, and TALE (transcription activator-like effectors) (16-20). For example, ZFP-Dnmts could specifically methylate whole *P16* CpG islands (8,16,17); TALE-Dnmts could methylate both target *P16* CpG islands and CpG islands within near regions (18); CRISPR/dCas9-Dnmts could focally methylate sgRNA target-flanking regions (50-bp, not including sgRNA target sequence) (21,22). Although the construction efficiency for ZFP-based editing tools is lower than those for dCas9- and TALE-based tools, it is likely that ZFP-Dnmts is optimal to induce full methylation of whole target CpG islands.

It has been reported that extensive methylation of whole *P16* CpG islands could be stably maintained in cancer cells, whereas focal methylation of *P16* exon-1 is not stable (23). In the present study, extensive methylation of whole *P16* CpG islands was stably and specifically induced in normal human fibroblasts by baseline P26-specific ZFP-Dnmt (P16-Dnmt). We found that *P16* methylation significantly prevented senescence, promoted proliferation, and prolonged lifespan of human fibroblasts.

## METHODS

### Cell line and Culture

The primary/early-passage normal human colon fibroblast CCD-I8C0 cell line was directly purchased from American Type Culture Collection (ATCC, CRL-1459) in March 2012. This cell line that was derived from normal human colon tissue from a 2.5-month old black female begins to senesce at subculture times 18 in our laboratory (the total population doubling time is about 42 according to ATCC specification description). Cells were cultured in ATCC-formulated EMEM medium (ATCC Cat. No. 30-2003) supplemented with 15% FBS (Gibco, USA), and maintained at 37°C in humidified air with 5 % C0_2_. Cells were subcultured with medium renewal every 2-3 days. When cell confluence reached 70-80%, cells were subcultured at ratio of 1:2-3. Cells subcultured at the 8^t^^h^ time in our laboratory (pre-experiment passage-8) were used in transfection experiments.

CCD-I8C0 cell line was tested and anthenticated by Genewiz, Inc Beijing on Sept. 9, 2014 when it was used in this study. Short tanden repeat (STR) patterns were analyzed using GenePrint 10 System (Promega). The data were analyzed using GeneMapper4.0 software and then compared with the ATCC databases for reference matching. This cell line is considered to be “identical” to the reference cell line CCD-I8C0 in the ATCC STR database, as the STR profile yields a 100% match.

### P16-Dnmt and Control Vectors and Transfection

The established P16-Dnmt and vector control pTRIPZ were prepared and used to transfect cells as previously described (8). Briefly, an engineered *P16* promoter-specific seven zinc finger protein (7ZFP) was fused with the catalytic domain of mouse *dnmt3a*, and integrated into the pTRIPZ vector containing a “Tet-On” switch. The purified P16-Dnmt plasmid DNA was mixed with pCMV-VSV-G and pCMV-Δ8.9 (Addgene, USA) to prepare lentivirus infection particles. When the confluence of CCD-I8C0 cells at pre-experiment passage-8 reached approximately 40%, these cells were infected with the P16-Dnmt or vector control virus particles, and incubated for 48 hours. Puromycin was added into the medium (final concentration, 0.5 |J.g/mL) to remove non-transfected cells, and to maintain the transfected cells. The pooled cells treated with puromycin for two weeks were considered to be stably transfected cells.

### Cell Proliferation Analysis using IncuCyte Zoom

Cell proliferation status was dynamically recorded using the IncuCyte ZOOM™ live-cell imaging platform. Each trial consisted of 6-9 parallel microwells, and all assays were repeated 2-3 times.

### Tumor Formation

Baseline P16-Dnmt or vector control CCD-I8C0 cells (5xl0^5^ cells/150 μiL) embedded with matrigel (BD Biosciences, NJ, USA) were subcutaneously injected (per site, 5 mice/group) into NOD-SCID mice (SPF-grade, female, 6-8 w, 20-22 g; purchased from Beijing HFK Biosci Co.LTD). Seventeen weeks after implantation, these mice were sacrificed, and the possible xenograft tumors were determined. The animal ethical committee at Peking University Cancer Hospital and Institute approved the experiment.

### Bisulfite-DHPLC and-Sequencing

Unmethylated cytosine residues in genomic DNA samples were modified with ZMYO EZ Methylation-God Kit (Cat# D5006). The 392-bp fragment within the *P16* exon-1 was amplified as previously described (24,25). The 468-bp fragment within the *P16* promoter was amplified with touchdown PCR (65°C to 50°C, −1°C/cycle) using a CpG-free primer set (forward, 5’-ggtgg ggttt ttata attag gaaag-3’; reverse, 5’-accct atccc tcaaa tcctc taaa-3’). The 392/468-bp PCR products were analyzed with DHPLC system (Transgenomics, USA) at the partial denaturing temperature of 58.6/55.5 ° C, and confirmed with clone sequencing.

### Genome-Wide Analysis of DNA methylation

Illumina Infinium HD Methylation 850K arrays were used to perform differential CpG methylation analyses on CCD-I8C0 cells stably transfected with the P16-Dnmt and pTRIPZ control vectors without doxycycline treatment according to the Assay Manual (by Capitalbio Technology Corporation, Beijing). DNA methylation ratio for probed CpG sites were computed as the ratio of normalized methylated signal intensity to the sum of methylated and unmethylated signal intensities using GenomeStudio software. Using the control vector at passage-5 as a reference,Δmethylation ratio was calculated to represent differential methylation for each CpG site in the baseline P16-Dnmt cells at passage-6, −9, (TSS-CGI) is more than 10% in the baseline P16-Dnmt cells at passage-6, and methylation ratio was consistently increased (at any level) in the baseline P16-Dnmt cells at passage-9 and -13, the differential Δ methylation was considered to be significant. When significant methylation was detected at 2 or more probed-CpG sites within a TSS-CGI, this CGI is defined as hypermethylated TSS-CGI. KEGG_PATHWAY enrichment was performed to analyze possible functions of differential methylation at probed CpG sites between cells with various treatments. The Methylation850K datasets were uploaded to NCBI with the accession number GSE111505.

### Western Blot and Confocal Analysis of P16 Expression Level

P16 protein levels in cells were analyzed as previously described (8). Rabbit monoclonal antibody against human P16 protein (abl08349, Abeam, Britain) was used in Western blot assay and mouse monoclonal antibody against human P16 protein (Ventana Roche-E6H4, USA) was used in the immunostaining assay.

### Detection of the Length of Telomere by Quantitative PCR

Copy number of the telomere gene was determined with quantitative real-time PCR (primer set: forward, 5’-cggtt tgttt gggtt tgggt ttggg tttgg gtttg ggtt-3’; reverse, 5’-ggctt gcctt accct taccc ttacc cttac cctta ccct-3’) as described (26). Non-telomere gene *36B4* was used as reference (primer set: forward, 5’-cagca agtgg gaagg tgtaa tcc-3’; reverse, 5’-cccat tctat catca aeggg tacaa-3’). The relative copy number was calculated using the formula (2’^,ΔCT^). Each experiment was performed in triplicate.

### Genome-Wide Analysis of Copy Number Variations

The genome-wide copy number variation was analyzed through construction of Library using TruSeq Library Construction Kit and sequenced with Illumina HiSeq XTen by Novogene (Beijing). The average sequencing depth was about 10 (Supplementary Dataset file 1

### β-Galactosidase Staining

**β**-Galactosidase *in situ* Staining Kit (GENMED, Shanghai) was used according to the User Manual. The experiments were repeated at least two times.

### Statistical Analysis

Results were displayed by constituent ratios of enumeration or ranked data. All P-values were two-sided, and a difference with P<0.05 was considered statistically significant.

## RESULTS AND DISCUSSION

Recently, we constructed a P26-specific DNA methyltransferase P16-Dnmt with seven-zinc finger protein (7ZFP) that could specifically bind to and induce full methylation of whole *P16* CpG islands in cancer cells (8). To study the effect of *P16* methylation on cancer development, a model of full methylation of *P16* CpG islands in normal diploid cells first needed to be established. Therefore, normal human primary CCD-I8C0 fibroblasts were stably transfected with P16-Dnmt. In pilot study, P16-Dnmt expression was induced by addition of doxycycline into the medium (dox; final concentration, 0.25 |J.g/mL). Three weeks after dox-treatment, a chromatographic peak for methylated-P26 alleles was observed in P16-Dnmt&dox fibroblasts in DHPLC analysis (Figure 1A). Such methylation peak was not detected in baseline P16-Dnmt fibroblasts (without dox-treatment), 7ZFP&dox, vector control&dox, and parental mock cells. Bisulfite-sequencing confirmed these results (Figure 1B). The entire *P16* CpG islands were extensively methylated in P16-Dnmt&dox cells with an average methylation density of 51.4%, and not methylated in vector control&dox cells. Interestingly, a few methylated CpG sites (6.8%) were also observed in baseline P16-Dnmt cells, indicating baseline P16-Dnmt expression in P16-Dnmt stably transfected fibroblasts without dox-treatment.

**Figure 1.**
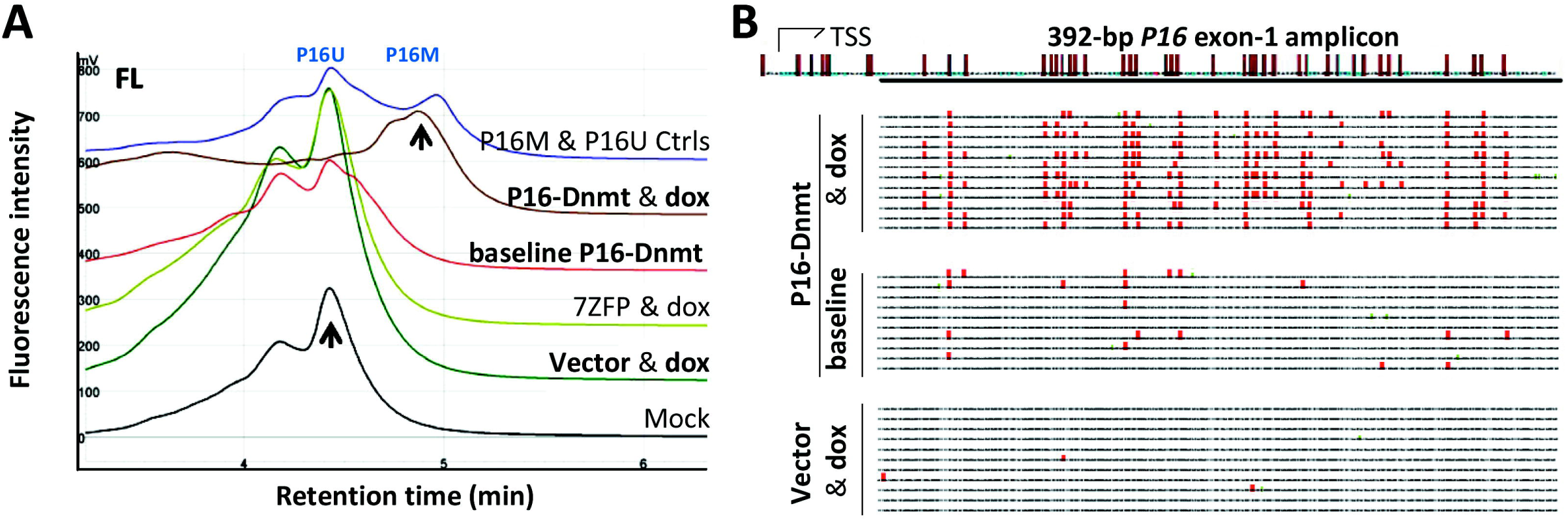
Induction of *P16* methylation in CCD-I8C0 fibroblasts by P16-Dnmt. **(A)** DHPLC chromatograms of methylated and unmethylated *P16* exon-1 PCR products (392-bp) for fibroblasts stably transfected with P16-Dnmt or vector control, with or without doxycycline treatment for 3 weeks. Genomic DNA sample from P26-hemimethylated HCT116 cells was used as methylated- and unmethylated-P26 reference controls (P16M & P16U Ctrls). **(B)** The results of bisulfite-sequencing of the 392-bp PCR products; each line represents a clone; red dots, methylated CpG sites; location of CpG sites within the amplicon was also displayed on the top. Methylation density was 51.4% (216/420) and 6.8% (19/280) for P16-Dnmt&dox and baseline P16-Dnmt cells, respectively.

It is well known that the *P16* gene plays a crucial role in cell senescence (27-29). To investigate whether *P16* methylation may prevent cell senescence, above pooled P16-Dnmt&dox, vector control&dox, and baseline P16-Dnmt control fibroblasts were continuously subcultured. Unexpectedly, both P16-Dnmt&dox and vector control&dox cells did not proliferate at passage-6 in the 10^th^ post-dox-treatment week, accompanied with cell size increase and cell disruption. As described blow, the same phenomenon was also observed in the baseline vector control cells without dox-treatment. In contrast, baseline P16-Dnmt fibroblasts remained to actively proliferate at this time point (Supplementary Figure 1). This suggests that baseline P16-Dnmt may be more efficient in preventing fibroblast senescence, probably due to less off-target effect or less cell stress in baseline P16-Dnmt cells.

To confirm whether baseline P16-Dnmt indeed prevents cell senescence and exclude the possible disturbance from dox-treatment, baseline P16-Dnmt fibroblasts and baseline vector control fibroblasts (without dox-treatment) were continuously observed using long-term IncuCyte live-cell imaging system with serial subcultures and medium renewals as described in the method section. The average proliferation rate of baseline P16-Dnmt cells (9 wells/group) was significantly higher than that of baseline vector control cells at the post-transfection subculturing number 5 (passage-5) (Figure 2, p<0.001). Notably, while baseline vector control fibroblasts stopped proliferating at passage-6, baseline P16-Dnmt cells continued to proliferate until passage-26 within the 53^th^ post-transfection week and did not become more abundant at later time points (Figure 3).

**Figure 2.**
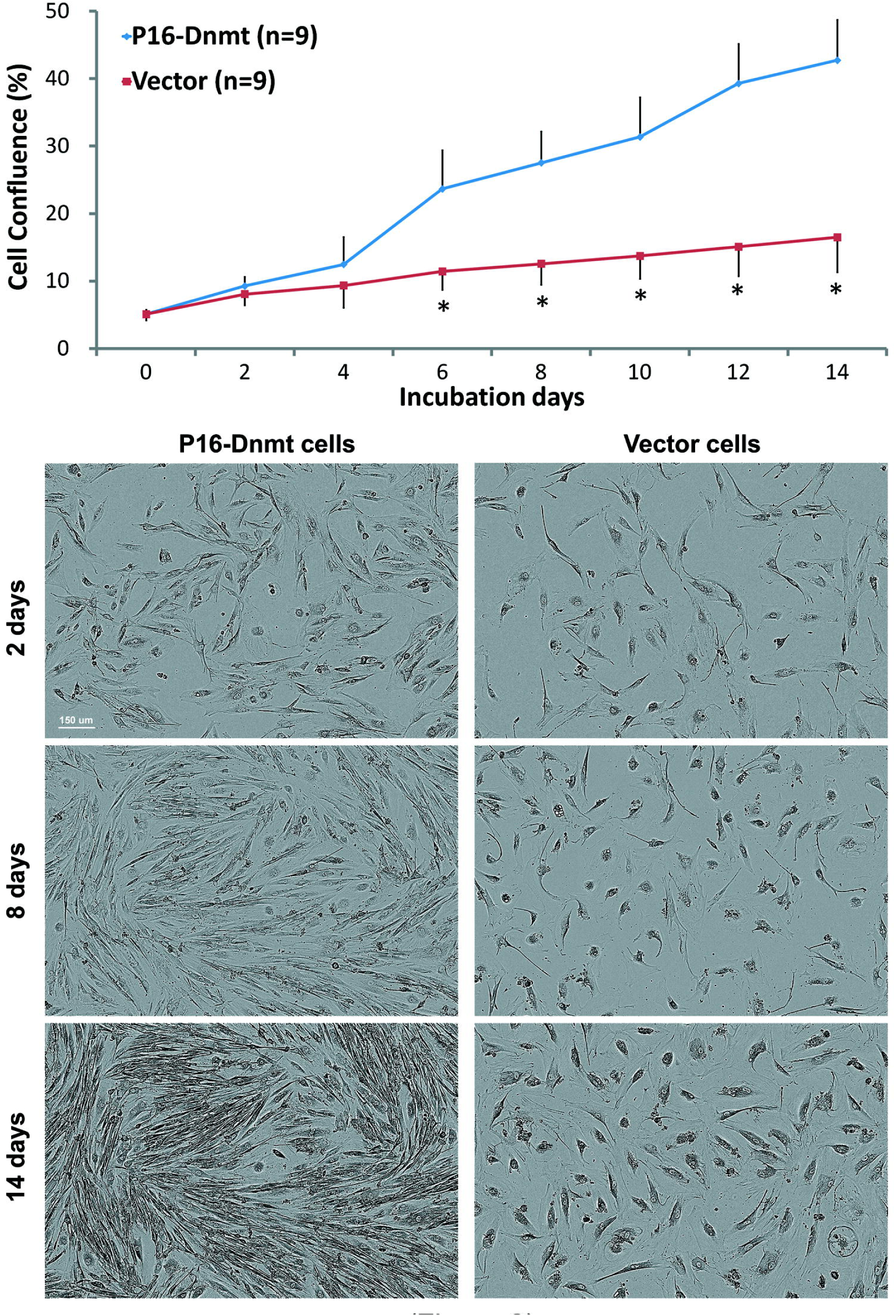
Proliferation curves for baseline P16-Dnmt and baseline vector CCD-I8C0 cells at passage-5 in IncuCyte Zoom analysis. Phase object images of live cells for two groups taken on three different post-subculture/incubation days were also listed, respectively.

**Figure 3.**
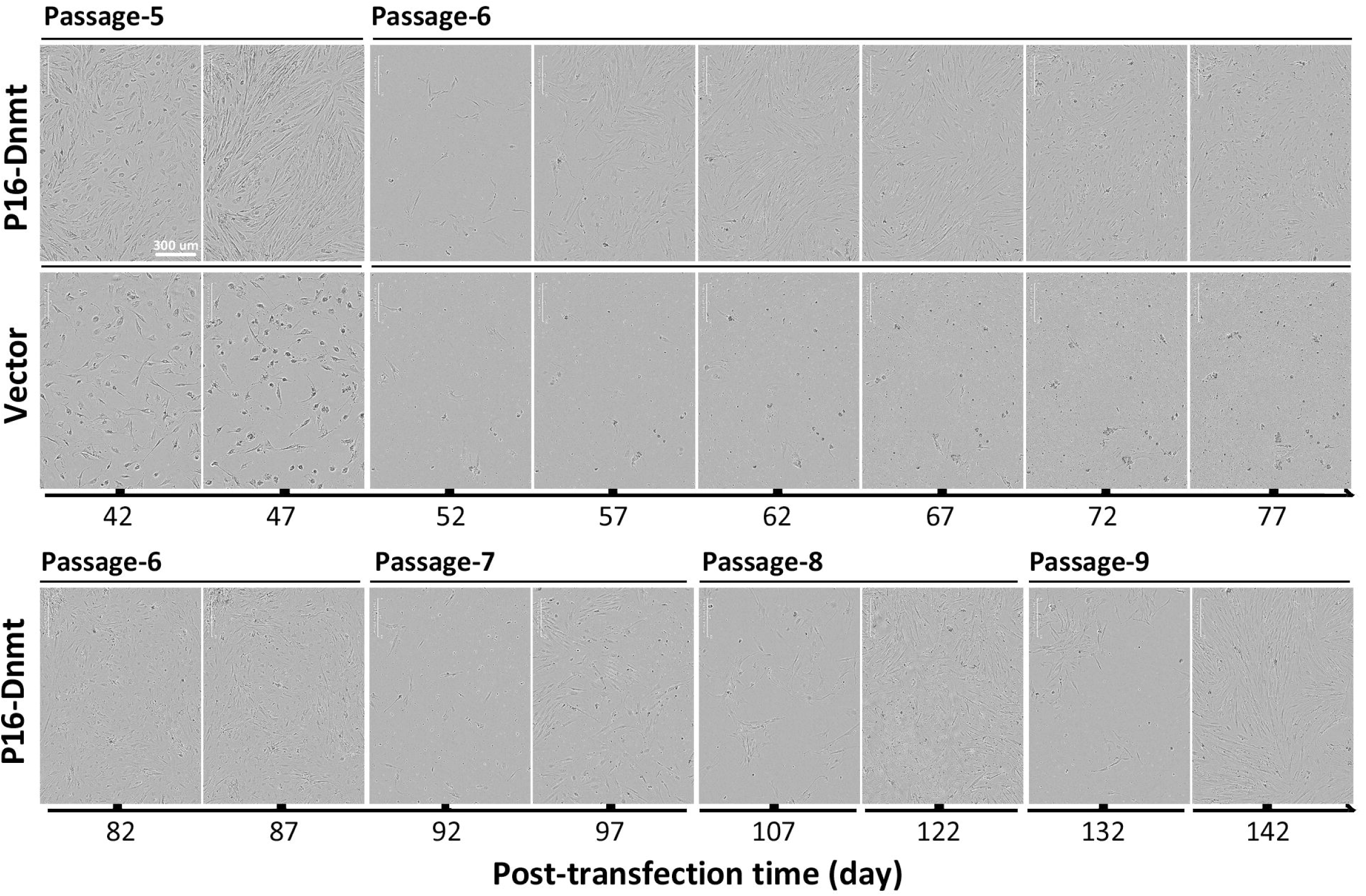
IncuCyte long-term observation of baseline P16-Dnmt and baseline vector control CCD-I8C0 cells. Both the post-transfection subculture day and passage number (subculture times) were labeled.

To evaluate the senescence status of these cells, β-galactosidase staining assay was used to detect senescing cells. The proportion of β-galactosidase-positive staining cells was significantly higher in baseline vector control cells than baseline P16-Dnmt cells at passage-5 (Figure 4, p<0.001). Together, these results indicate that baseline P16-Dnmt expression could promote proliferation of human primary fibroblasts and prevent cell senescence.

**Figure 4.**
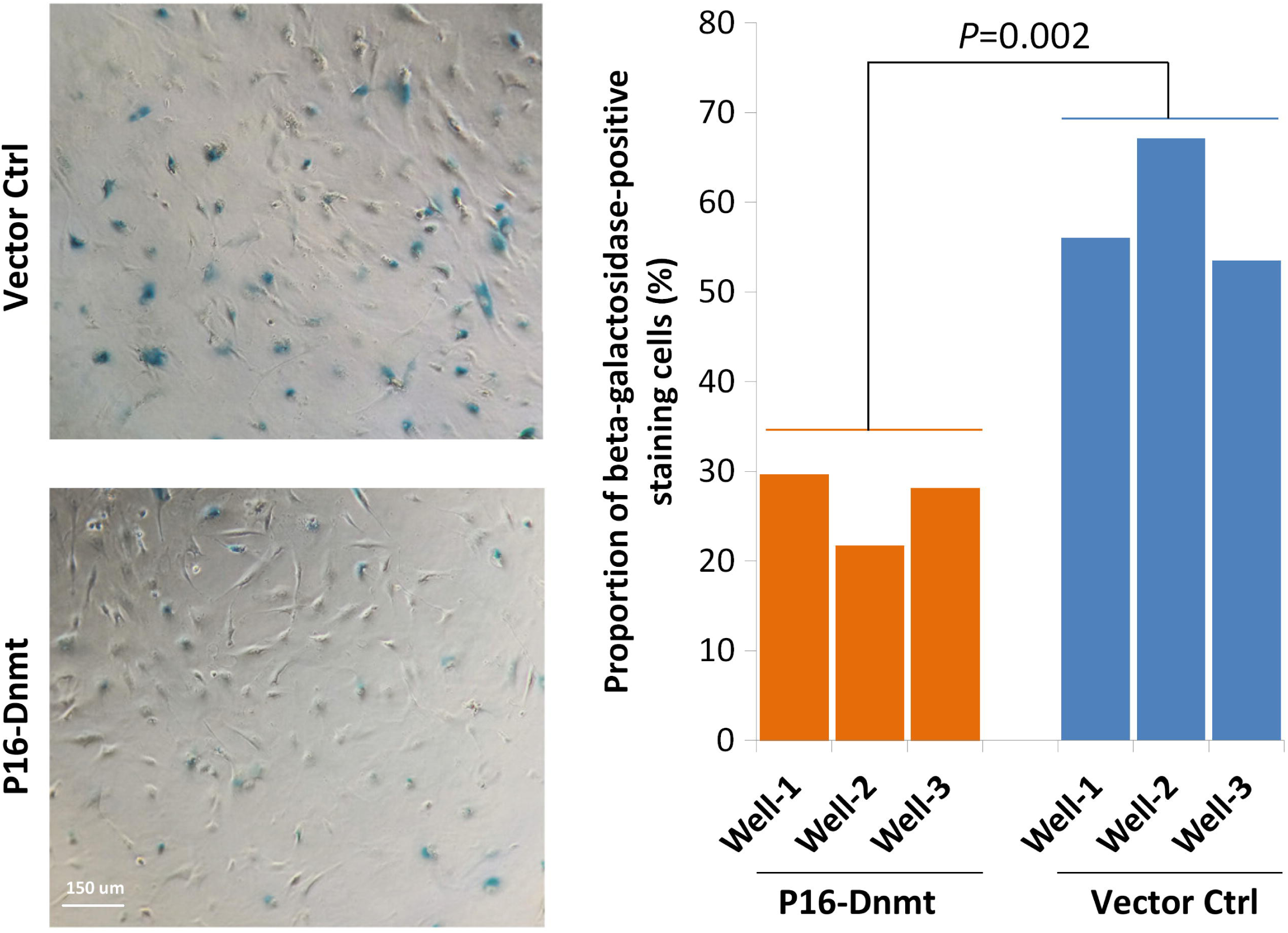
Comparison of the images and proportion of β-galactosidase-positive staining cells in baseline P16-Dnmt cells and baseline vector control cells at passage-5. Student t-test was used to calculate the P-value between the average ratio of β-galactosidase-positive staining cells to total CCD-I8C0 fibroblasts between baseline P16-Dnmt cells and baseline vector control cells in triplicate.

To study whether baseline P16-Dnmt indeed induce *P16* methylation in these fibroblasts, the methylation status of *P16* alleles in these cells were dynamically analyzed using Illumina Methylation850K array. The prevalence of methylated CpG sites were gradually increased at the probed CpG sites within entire *P16* CpG islands in baseline P16-Dnmt CCD-I8C0 cells along with cell subculture (Figure 5A, red-dot-line squared). Such phenomenon was not observed in *P14* and *P15* CpG islands located within the same gene locus (Supplementary Figure 2A). Although hypermethylation was also found in most CpG sites within the *MMP28* CpG islands, hypermethylation at only sporadic CpG sites were observed within non-target CpG islands, including *CCNE2, CDH2, GATA5, RUNX3, TERT*, and *WT1* genes (Supplementary Figure 2B). Because P16-Dnmt could methylate entire target CpG islands, thus, these sporadic CpGs may not be directly methylated by P16-Dnmt. Instead, hypermethylation at these sporadic CpGs may be one kind of adaptive response in fibroblasts escaping from senescence.

**Figure 5.**
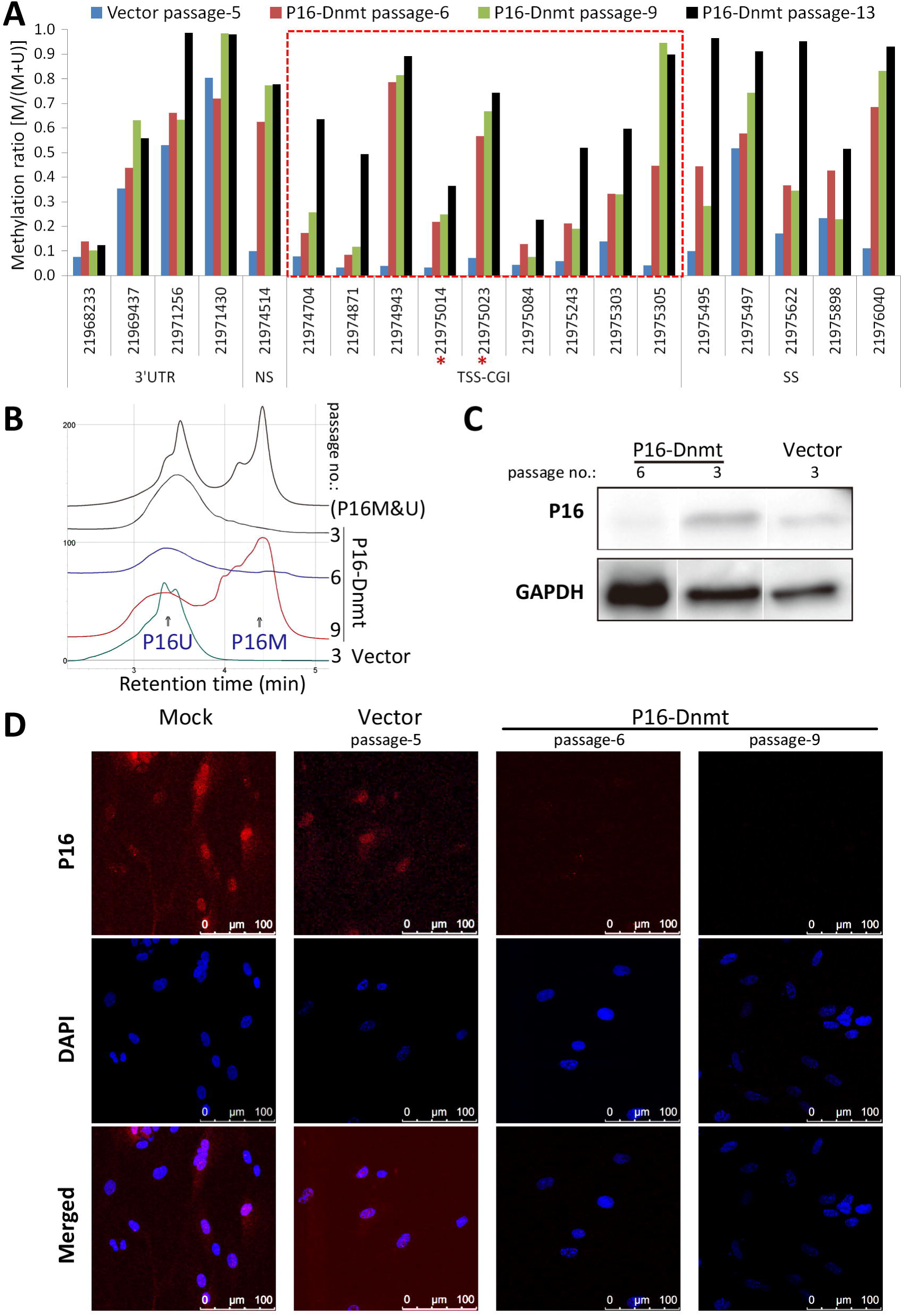
Effects of P16-Dnmt expression at baseline level on the methylation and expression status *of P16* alleles in CCD-I8C0 cells at different passage number. **(A)** Methylation status of each lllumina 850K methylation array probed-CpG sites within various *P16* gene regions, including N-shore (NS) and S-shore (SS), in baseline vector control cells (passage-5) and baseline P16-Dnmt cells (passage-6/9/13; *, CpG site with significant differential methylation). **(B)** DHPLC chromatograms of methylated and unmethylated *P16* promoter PCR products (468-bp) for fibroblasts stably transfected with P16-Dnmt or vector control. **(C)** Results of Western blot analysis for detecting the P16 protein (three lanes reorganized from the same original image). **(D)** Results of confocal immunostaining for detecting the P16 protein *in situ.*

The DHPLC results further showed that the methylated-P16 proportion was gradually increased in baseline P16-Dnmt cells along with cell subcultures (0 at passage-3, 20.2% at passage-6 and 68.8% at passage-9; Figure 5B). Bisulfite-sequencing confirmed the Methylation850k and DHPLC analyses. The average methylation density was 28.1% and 41.6% at passage-6 and passage-9, respectively, and further increased to 63.2% at passage-13 within the 468-bp promoter CpG islands (Supplementary Figure 3A). Similar results were also observed within *P16* exon-1 region (Supplementary Figure 3B). The above results indicate that baseline P16-Dnmt could efficiently and specifically induce *P16* methylation in CCD-I8C0 cells.

The results of Western blot analysis in this study showed that *P16* expression was significantly lost in baseline P16-Dnmt cells at passage-6 (Figure 5C). In the confocal immuno-fluorescence analysis, no P16-positve cell was observed in baseline P16-Dnmt cells at passage-6 and passage-9, while P16 protein was observed in the nucleus of parental mock and baseline vector control fibroblasts at passage-5 (Figure 5D). Generally *P16* expression is repressed in embryonic/tissue stem cells, induced-pluripotent stem cells (iPSC), and many immortalizing cells (30-33). Our results indicate that *P16* expression is comprehensively inactivated in baseline P16-Dnmt cells, which prolongs the lifespan of fibroblast cells, reminiscent of the silencing of *P16* in the aforementioned cells.

To explore possible reason for the behavior differences between baseline P16-Dnmt and vector control cells, the length of telomeres was determined with quantitative PCR. The results unveiled that the average telomere length for baseline P16-Dnmt cells gradually decreased along with cell subculture (66.4% and 24.5% at passage-4 and passage-5 relative to vector control at passage-4; Supplementary Figure 4). Interestingly, the KEGG_PATHWAY analysis results showed that top 3 pathways enriched with most differential methylation at Methylation850k dataset between vector control cells at passage-5 and baseline P16-Dnmt cells at passage-6 were metabolic pathways, pathways in cancer, and PI3K-Akt signal pathways (Figure 6A); between baseline P16-Dnmt cells at passage-6 and passage-13, PI3K-Akt signal pathways, pathways in cancer, and metabolic pathways (Figure 6B).

**Figure 6.**
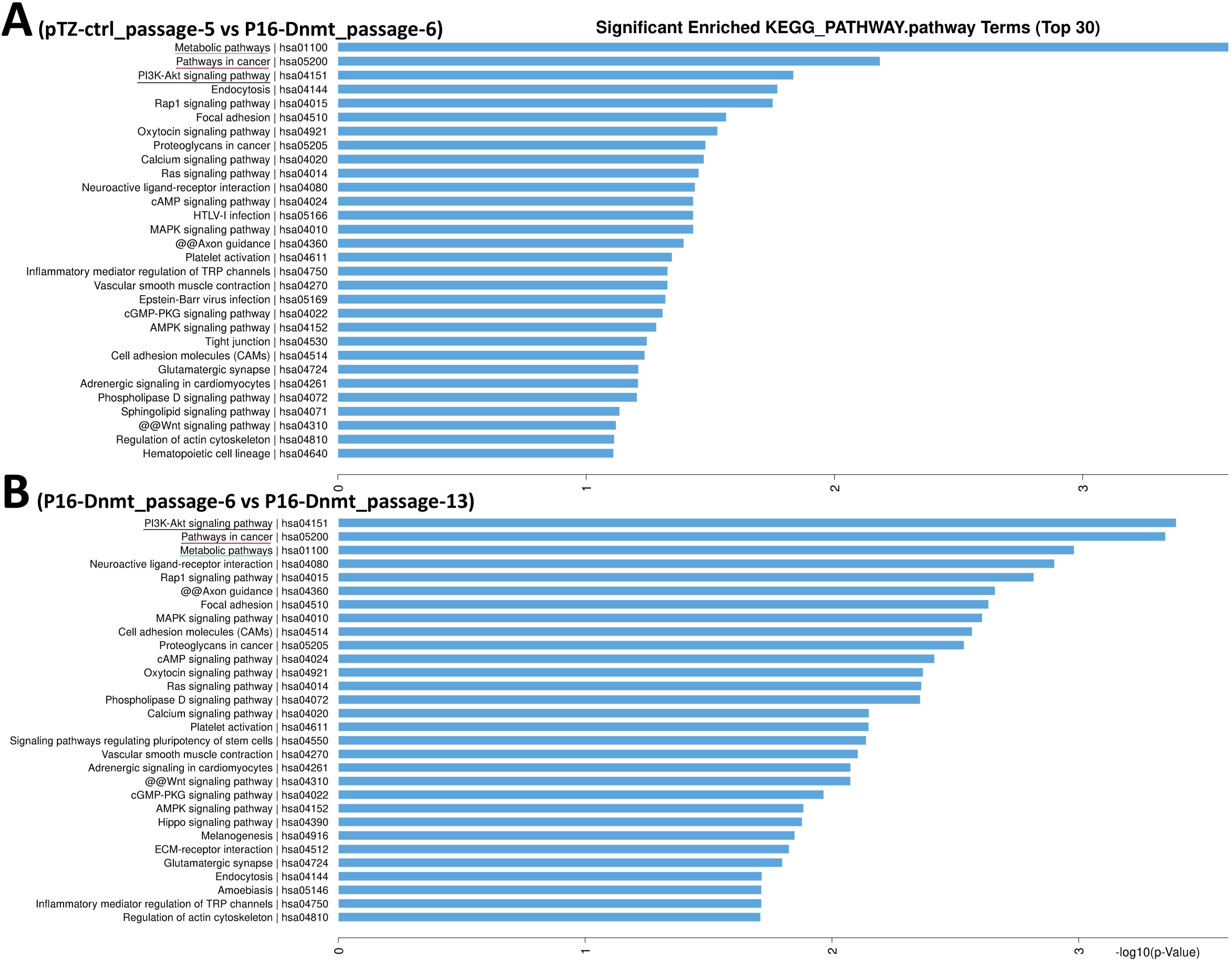
Top 30 KEGG_PATHWAY pathways enriched with differential methylation at probed-CpG sites in lllumina 850K methylation analysis between CCD-C0I8 fibroblasts with and without baseline P16-Dnmt treatment. **(A)** between pTZ vector control cells at passage-5 and baseline P16-Dnmt cells at passage-6; (B) between baseline P16-Dnmt cells at passage-6 and passage-13.

We further investigate whether baseline P16-Dnmt or vector control cells can grow xenograft in NOD-SCID mice. As expected, no xenograft was formed in NOD-SCID mice subcutaneously injected both cells coated with Matrigel at the 17^th^ post-implantation week. A similar finding was also reported by others using TALE- or CRISPR/dCAS9-based tool to induce partial *P16* methylation (18,21). The occurrence of gradually shorten telomere length in baseline P16-Dnmt cells and the inability to form xenograft in mouse may account for the idea that *P16* methylation could prevent the senescence and expand the lifespan, but could not immortalize human fibroblasts. However, oncogenic stimuli can induce oncogene-induced senescence in fibroblasts to protect against transformation, and *P16* inactivation/downregulation can prevent oncogene-induced cell senescence and thus sensitize cells to malignant transformation (34-37). It would be interesting to determine whether epigenetic *P16* inactivation may cooperate with other tumor-related genes or endogenous/environmental carcinogens to affect cancer development.

It was reported that *P16* could prevent centrosome dysfunction and genomic instability in primary cells and that shRNA-knockdown of *P16* expression generated supernumerary centrosomes through centriole pair splitting in human diploid epithelial cells (38-40). To study whether *P16* inactivation by DNA methylation affects genome stability of primary diploid fibroblasts, we analyzed the whole genome copy number changes with next-generation-sequencing. No detectable chromosome/gene copy number variation was detected between the genomes of baseline P16-Dnmt and vector control cells at passage-4 (Supplementary Dataset file 1).

Gene expression-controllable vectors such as pTRIPZ vector containing a “Tet-On” switch are often used in transfection studies to avoid uncontrollable gene overexpression. However, gene baseline expression in pTRIPZ vectors is unavoidable in cells even without dox-treatment (41-43). We found that only baseline P16-Dnmt cells could escape from senescing at the same time point at which P16-Dnmt&dox and vector control cells stopped proliferating. That 68.8% *P16* alleles were methylated in these cells indicates that the occurrence of baseline P16-Dnmt expression could lead to subsequent on-target DNA methylation. This also implies that gene baseline expression in pTRIPZ vectors might be a good strategy for gene function studies.

In conclusion, the present study unveils that *P16* methylation could prevent cell senescence, promote cell proliferation, and expand the life span of human fibroblasts, which may contribute to cancer development.

*Conflict of Interest Statement:* None declared.

## Abbreviations

DHPLC: denatured high performance liquid chromatography
Dnmt: DNA methyltransferase
P16-Dnmt: P26-specific ZFP-Dnmt
TSS: transcription start site
ZFP: zinc finger protein

## Acknowledgements

This work was supported by Beijing Municipal Science and Technology (#Z151100001615022) and the National Natural Science Foundation of China (#81773036 and #91640108) and the National Basic Research Program of China (973 #2015CB553902). We appreciate Kendra Williams (USA) for English language editing.

## Author contribution

YG performed the cell culture experiment. CC constructed the P16-Dnmt vectors. SX constructed vector for seven-zinc finger protein. BZ analyzed the methylation pattern. DD designed this study, analyzed the data, and wrote the manuscript. All authors have read and commented on the manuscript and approved the final version.

## Supplementary data

**Supplementary Figure 1.** Morphology of CCD-I8C0 fibroblasts stably transfected with P16-Dnmt or empty control vector, with or without doxycycline treatment at the 3^rd^ and 9^th^ post-doxycycline treatment week. Giemsa’s *in situ* staining.

**Supplementary Figure 2.** Methylation ratios of probed CpG sites within baseline P16-Dnmt on-target and off-target CpG islands in baseline P16-Dnmt (at passage-6/9/13) and vector control CCD-I8C0 reference (at passage-5) in Illumina Methylation 850K array analysis. The methylation status of CpG sites in CpG islands around transcription start site (TSS-CGI) and other regions within *P16, P14* and *P15* genes **(A)** and other TSS-CGIs **(B).** NS, N-shore; SS, S-shore; NF, N-shelf; SF, S-shelf. Red mark (*), significantly hypermethylated probed-CpG site; Red line squared CpG sites, CpG sites within the TSS-CGI

**Supplementary Figure 3.** Bisulfite-sequencing for the 468-bp and 392-bp PCR products derived from the *P16* Promoter **(A)** and Exon-1 **(B)** in CCD-I8C0 fibroblasts stably transfected with P16-Dnmt or empty control vector without doxycycline treatment at different passages. The listed average methylation density value was calculated according to ratio of actual methylated CpG sites to the total CpG methylation candidates within the informative clones in each group. The consensus sequences of these amplicons are illustrated on the top-lines. Each red dot represents a methylated CpG site. Each line represents a clone.

**Supplementary Figure 4.** Comparison of the average length of telomere in baseline P16-Dnmt and control CCD-I8C0 cells. Student t-test was used to calculate the P-value between the average telomere lengths for various CCD-I8C0 fibroblast groups (3 wells/group).

**Supplementary Dataset file** 1. Genome-wide copy number analysis of CCD-I8C0 cells stably transfected with P16-Dnmt and control vector, with and without doxycycline treatment.

